# High Throughput-based Surveillance Reveals New Components of Spanish Citrus Virome Related to Tristeza, Impietratura and Yellow Vein Clearing Diseases

**DOI:** 10.64898/2026.02.12.705513

**Authors:** María Martínez-Solsona, Luis Felipe Arias-Giraldo, Antonio Olmos, Félix Morán, Ana Belén Ruiz-García

## Abstract

Citrus is one of the most important crops cultivated worldwide, representing a strategic source of agricultural income for many countries, including Spain, the main producer within the European Union. The protection of this highly valuable industry against an increasing global movement of pests and pathogens requires effective regulatory measures, including control of plant propagation material, phytosanitary surveillance and risk assessment, which are based not only on knowledge of the established threats but also on potential emerging threats affecting citrus that may circulate unnoticed in the production system. In this work, with the aim of generating knowledge on potential emerging viruses in Spanish citrus orchards, high-throughput sequencing (HTS) analysis has been applied to monitor the sanitary status of several growing areas of one of the main citrus producer regions in Spain, the Valencian Community. The results of this study have revealed a much more complex citrus virome than previously reported, including citrus yellow vein clearing virus (CYVCV), a non-regulated but harmful citrus virus, as well as the T3 genotype of citrus tristeza virus (CTV) and citrus virus A (CiVA), not detected to date in Spain. Moreover, our results indicate the existence of other unknown components of the citrus virome. HTS detection of CYVCV, CTV T3 and CiVA and their presence in Spanish orchards has been confirmed by RT-PCR and Sanger sequencing. These findings have relevant implications in the development of control and regulatory measures against three important viral diseases, tristeza, impietratura and yellow vein clearing diseases, and demonstrate the added value of HTS-based surveillance to discover emerging components of the citrus virome.

---

The citrus industry in Spain is essential to the agricultural sector, recognized for its strong export orientation and high-quality standards. Spain is the largest citrus exporter and the main producer within the European Union (EU) with 294,680 cultivated hectares and a total citrus production of 5,727,280 tons according to FAOSTAT (2024). Among Spanish citrus-growing regions, the Valencian Community contributes with almost half of the national production, followed by Andalucía and Murcia. Despite this success and the existence of a centralized quarantine, sanitation and certification framework, that has played a key role in the renewal, maintenance and reinforcement of citrus industry, Spanish citriculture faces significant challenges from diseases that threaten fruit yield and quality. Climate change, the spread of insect vectors, and the intensification of plant movement are increasing the risk of emerging pests and pathogens (Singh *et al*., 2023). In this scenario, the emergence of new viral diseases represents one of the most critical constraints for the sustainability of citrus production. Consequently, production systems must be continuously updated to support advanced diagnostics, surveillance, and regulatory schemes to effectively respond to the real and evolving complexity of viral threats affecting citrus.

Many viruses have been described to infect citrus crops, with some of them responsible for important diseases that have a significant impact on fruit production (Zhou *et al*., 2020). According to their potential phytosanitary impact and geographical distribution in the EU, citrus infecting viruses can be broadly categorized in three regulatory groups: quarantine viruses, regulated viruses and non-regulated viruses, in accordance with Regulation (EU) 2016/2031 and its implementing Regulation (EU) 2019/2072 on protective measures against plant pests (European Commission, 2019). Quarantine viruses constitute the most severe threat due to their potential to cause devastating economic losses and their restricted or absent distribution within the EU. This group includes non-EU genotypes of citrus tristeza virus (CTV) (EFSA PLH Panel *et al*., 2017), satsuma dwarf virus (SDV) and the citrus leprosis-associated viruses (CLaVs) caused by viruses belonging to two distinct genera, *Cilevirus* (citrus leprosis virus C, CiLV-C; and citrus leprosis virus C2, CiLV-C2) and *Dichorhavirus* (citrus strain of orchid fleck virus, OFV-citrus; citrus leprosis virus N, CiLV-N; and citrus chlorotic spot virus C, CiCSV) (Zhou *et al*., 2020). Regulated non-quarantine viruses include pathogens that, although already present, are subject to phytosanitary measures due to their impact on production and trade. However, according to the current EU phytosanitary legislation, only a limited number of citrus viruses and viroids are regulated. These include two viroids, citrus exocortis viroid (CEVd) and hop stunt viroid (HSVd); four viruses, citrus psorosis virus (CPsV), citrus variegation virus (CVV), citrus leaf blotch virus (CLBV) and EU genotypes of CTV; as well as two additional entities, the citrus cristacortis agent and the citrus impietratura agent (European Commission, 2019; Zhou *et al*., 2020). The two latter are listed in the regulation as “agents” because their etiological causes have not been conclusively demonstrated, although recent studies indicate that impietratura disease is associated with a recently identified virus named citrus virus A (CiVA) (Beris *et al*., 2021).

In contrast to the limited number of regulated pathogens, a much larger set of citrus viruses and viroids with proven or potential phytosanitary relevance remains outside the current regulatory framework. Several of these agents have been associated with severe diseases and significant economic losses in citrus-growing regions worldwide. Several citrus viroids remain unregulated, citrus bent leaf viroid (CBLVd), hop stunt viroid (HSVd) in citrus, citrus dwarfing viroid (CDVd), citrus bark cracking viroid (CBCVd), citrus viroid V (CVd-V), citrus viroid VI (CVd-VI) and citrus viroid VII (CVd-VII) and other important viruses as well including citrus vein enation virus (CVEV), citrus tatter leaf virus (CTLV), citrus chlorotic dwarf-associated virus (CCDaV), Indian citrus ringspot virus (ICRSV), citrus mosaic virus (CYMV), citrus sudden death-associated virus (CSDaV) and citrus yellow vein clearing virus (CYVCV) (Zhou *et al*., 2020). This latter one is included in the EPPO alert list although it is not included in the Regulation (EU) 2016/2031 and its implementing Regulation (EU) 2019/2072. In addition, the continuous identification of new viruses and viroids as well as the scarce knowledge on their potential impact, could lead to important gaps in current regulation systems that would have to be covered in order to successfully address the full diversity of emerging phytosanitary threats.

High-throughput sequencing (HTS) has become a key tool in plant virology, significantly advancing our understanding of viral diversity. The continuous development of sequencing platforms, together with increasing capacity and more affordable costs, has made HTS accessible for many research and diagnostic applications. Thus, its relevance goes beyond traditional diagnostic methods. In perennial crops such as citrus, mixed infections are common and symptom expression is often variable or absent, which limits the performance of surveillance strategies based exclusively on targeted diagnostics (Chen *et al*., 2025). As a result, a proportion of the viral diversity present in orchards and propagation material may remain undetected when surveys focus only on a predefined list of regulated pathogens. This situation is particularly relevant in production systems that rely on certification schemes, where undetected infections may contribute to the dissemination of emerging or poorly characterized agents. From a phytosanitary perspective, HTS provides information that is difficult to obtain with conventional diagnostic tools and supports a more realistic assessment of phytosanitary risk. Virome-level analyses allow the simultaneous detection of regulated and non-regulated agents in the same assay, facilitate the identification of newly introduced viruses and genotypes, and provide a framework to monitor changes in virus populations over time. In a regulatory context in which the list of targeted pathogens is necessarily limited and periodically updated, HTS-based surveys can help to identify gaps in current surveillance systems and to prioritize agents that may require further biological characterization and/or risk assessment. In this sense, the integration of HTS into citrus surveillance programs represents a valuable complement to existing diagnostic and regulatory strategies, particularly in scenarios characterized by intense plant movement, high vector pressure and rapid epidemiological change.

In this context, the aim of the present study was to characterize the citrus virome in representative citrus-growing areas of one of the main citrus producing areas in Spain, The Valencian Community, using HTS-based approaches, with a focus on the detection of regulated pathogens, emerging viruses and previously unreported viruses. The analysis of 20 datasets from orchards in this region led to the identification of several relevant viral agents, including regulated viruses, non-regulated viruses with recognized phytosanitary relevance such as CYVCV, and candidate novel citrus viruses. Importantly, recent EPPO initiatives to evaluated CYVCV within Pest Risk Analysis (PRA) schemes reflect the growing recognition of its phytosanitary relevance.

The study further aimed to assess the added value of virome-level analyses for phytosanitary surveillance and to provide data that may support future updates of surveillance strategies and regulatory priorities in Spanish and Mediterranean citrus production systems.

## Material and Methods

### Plant material and sample preparation

In an initial survey conducted in the Valencian Community covering the three provinces of Valencia, Castellón, and Alicante a total of 20 citrus samples from 13 different orchards were collected. The sampling included 11 *Citrus clementina* trees and 9 *Citrus sinensis* trees. These citrus plants were in the province of Valencia (16 samples, 8 orchards), Castellón (3 samples, 3 orchards), and Alicante (1 sample, 1 orchard). Samples represented the most common cultivars and rootstocks (Supplementary table S1) and included both asymptomatic and symptomatic plants showing tree decline, leaf deformation and stunting. Based on the HTS results obtained from this initial survey, a second sampling was carried out in citrus orchards located in the same orchard and/or in closely located orchards where viruses of agronomic relevance were detected. This survey comprised 68 additional citrus samples collected in the provinces of Castellón, Valencia and Alicante. The citrus species and cultivars included in this second sampling are detailed in Supplementary table S2.

For each plant, young leaves were collected individually in plastic bags (Bioreba, Reinach, Switzerland) and transported to the laboratory under refrigerated conditions. Samples were homogenized in extraction buffer (PBS containing 0.2% diethyldithiocarbamate and 2% PVP-10) at a 1:5 (w:v) ratio. Homogenization was performed using a Homex 6 homogenizer (Bioreba, Reinach, Switzerland) and resulting plant extracts were kept on ice until subsequent processing.

Total RNA was purified from 200 µL of plant extract using the Plant/Fungi RNA Isolation Kit (Norgen Biotek Corporation, Thorold, ON, Canada) following the manufacturer’s instructions, including the DNase treatment step, performed with RNase-Free DNase I (Norgen Biotek Corporation, Thorold, ON, Canada). Total RNA concentration was determined by spectrophotometry using a DeNovix DS-11 spectrophotometer (DeNovix Inc., Wilmington, DE, USA) and RNA samples stored at -80 °C until further analysis.

### High-Throughput Sequencing (HTS)

Total RNA integrity was evaluated using the Agilent 4200 TapeStation System (Agilent, Part# G2991BA). Libraries were prepared with the TruSeq Stranded Total RNA with Ribo-Zero kit (Ilumina, CA, USA) according to the manufacturer’s instructions, Prep Guide, Part #15031048. Paired-end sequencing (2 x 150 bp) was performed on an Illumina NovaSeq X platform (Ilumina, CA, USA). Sequencing quality was assessed using raw data statistics, including total reads, Q20 and Q30 (%). Sequencing runs were considered successful when at least 15 million paired-end reads per sample were obtained, Q20 values exceeded 99%, and Q30 values were above 90%.

### Bioinformatic preprocessing

HTS raw data were subjected to the next bioinformatic workflow analyses: initial adapter trimming and read quality processing were performed with fastp v1.0.1 (Chen *et al.,* 2023). Quality assessment was summarized with MultiQC (Ewels *et al*. 2016) before and after the trimming step. Taxonomic profiling of pre-processed reads was performed using Kraken2 v.2.17.1 (Wood *et al.,* 2019) with PlusPFP v.10152025 (https://benlangmead.github.io/aws-indexes/k2), and the results were visualized with recentrifuge v.1.16.1 (Martí 2019). Unclassified reads and those classified as viral were retained with krakentools v.1.2.1 (Lu et al., 2022). Remaining rRNA reads were removed with RiboDetector v.0.3.2 (Deng *et al*., 2022).

*De novo* assembly of viral sequences was performed with SPAdes v4.2.0 in rRNA mode (configured with flags -k 21, 33, 55, 77, 99, 127 and --only-assembler) (Bushmanova *et al*., 2019). Contigs longer than 200 bp were retained for subsequent analysis. geNomad v1.11.2 (Camargo *et al*., 2023) was used in relaxed mode (--relaxed) to identify, annotate, and taxonomically classify viruses and other mobile elements; subsequently, CheckV v1.0.3 (Nayfach *et al*., 2021) was used to estimate genome completeness of contigs, classifying them into five categories: complete, high quality (>90% completeness), medium quality (50–90% completeness), low quality (0–50% completeness), or undetermined quality (no completeness estimate available). Finally, BLASTn from the BLAST+ package v2.9.0 (Altschul *et al*., 1990) was used to generate local alignments between all assembled contigs and the Reference Viral DataBase v31.0 using the non-clustered sequences (Chin *et al*., 2025; Goodacre *et al*., 2018).

Contigs classified as high quality by CheckV were selected for downstream analyses. Selected contigs were used as references for read mapping-based extension using Geneious Prime software 2026 (Biomatters Ltd., Auckland, New Zealand), allowing the recovery of near full-length viral genomes. In parallel, the same set of contigs were characterized by BLASTX searches against the NCBI database “clustered nr” and by conserved domain analysis using the NCBI Conserved Domain Database (CDD v3.21) (Wang J *et al*., 2023), enabling the identification of hallmark viral proteins and conserved functional motifs. To identify viroid-like sequences, all contigs ranging from 150 to 500 nt in length were screened by BLASTn against a custom RefSeq-based database comprising representative sequences of the families *Avsunviroidae* and *Pospiviroidae*.

### Virus Detection and Confirmation by RT-PCR and Sanger Sequencing

CYVCV, CiVA, and CTV T3 were tested by RT-PCR using newly designed primers or primers previously reported in the literature, modified based on the HTS sequences recovered in this study (Table 1).

**Table 1.**
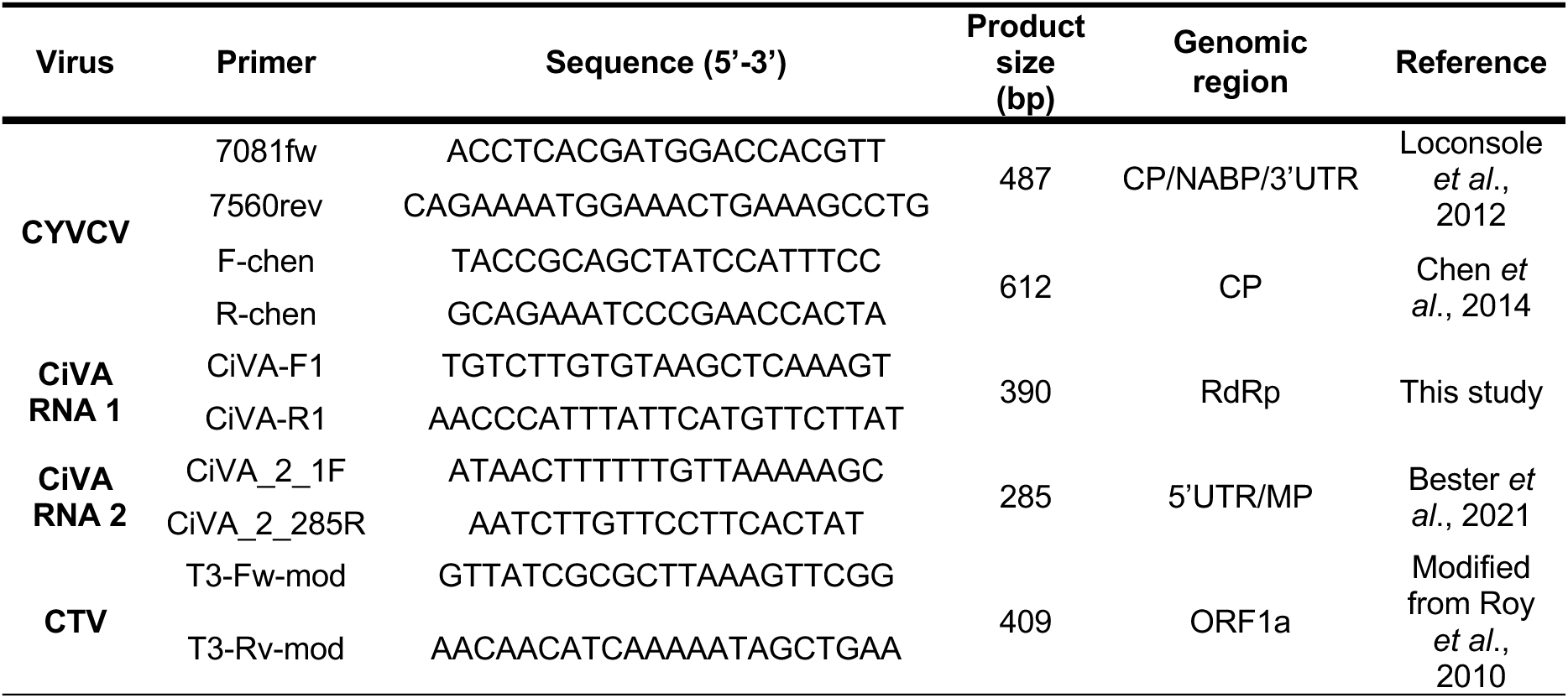
Primers used for RT-PCR detection of CYVCV, CiVA, and CTV T3. The table lists the target virus, primer sequences, expected amplicon size, its genomic region and the corresponding reference.

Primer pairs for the identification of CiVA RNA1 were designed based on the HTS sequence recovered in this study (PX970363). For the detection of CTV T3, primers from Cook *et al*. (2016) were modified based on the sequences recovered in this study (PX970361 and PX970362). Primer candidates were evaluated in silico using the OligoAnalyzer Tool (IDT) to assess melting temperature, GC content and secondary structures.

All RT-PCR reactions were carried out using AgPath One-Step RT-PCR kit (Applied biosystems, Foster City, CA, USA) following the manufacturer’s instructions. The reaction mixture contained 0.5 µM concentration of each primer and 2 µl of total RNA in a final volume of 25 µL. Amplification protocol consisted of: for the two pairs of primers of CYVCV, one step at 45°C for 45 min, one step at 95°C for 10 min and 35 cycles of amplification (95°C for 15 s, 55°C for 30 s, and 60°C for 45 s), with a final step at 60°C for 10 min; for CTV T3 and CiVA RNA1, one step at 45°C for 45 min, one step at 95°C for 10 min, and 35 cycles of amplification (95°C for 15 s, 55°C for 30 s, and 60°C for 1 min), with a final step at 60°C for 10 min; and for CiVA RNA2, one step at 45°C for 45 min, one step at 95°C for 10 min, and 35 cycles of amplification (95°C for 15 s, 50°C for 30 s, and 60°C for 1min 30 s), with a final step at 60°C for 10 min.

Amplicons were Sanger sequenced in both directions after purification using mi-PCR Purification Kit (Metabion International AG, Martinsried, Germany) following the manufacturer’s instructions.

### Phylogenetic Analysis

For CYVCV, a total of 80 complete genome sequences representing isolates from different geographic regions, including Asia, Europe and the USA, were retrieved from GenBank and included in the analysis. For CiVA, a total of 21 complete RNA1 sequences retrieved from GenBank were used, representing isolates from Europe (Italy, France, and Greece), Africa (South Africa), Asia (China, India, and Turkmenistan), Australia and the USA. For CTV, a total of 141 complete genome sequences representing the major phylogenetic groups and biological variants were retrieved from GenBank and included in the analysis. The dataset comprised isolates from Asia, Europe, Africa, the Americas, and Oceania, encompassing mild and severe strains associated with stem pitting and decline, including representative genotypes such as VT, T30, T36, T3, T68, RB, HA16-5, and S1. All multiple sequence alignments were generated using MAFFT implemented in Geneious Prime 2026 software (Biomatters Ltd., Auckland, New Zealand) and manually inspected to ensure positional homology. The best substitution model for the phylogenetic analysis of CYVCV and CiVA was obtained using MEGA v12.1 software (Stecher *et al*., 2025). General time-reversible (GTR) with a discrete gamma distribution to account for evolutionary rate differences among sites (+G) and allowing for a proportion of invariant sites (+I) was used in the two cases. Phylogenetic trees were constructed with the maximum likelihood method and branch support was assessed using 1000 bootstrap replicates. Trees were inferenced CYVCV and CiVA by IQ-TREE 3 (Wong et al., 2025) and MEGA v12.1, respectively. In the case of CTV, an unrooted phylogenetic network was constructed using SplitsTree4 v4.16.2 to identify the different genotype groups (Bester *et al*.,2020). Accession numbers, isolate names, and geographic origins are indicated directly in the phylogenetic trees, and the corresponding GenBank records are cited in the reference list.

## Results

### HTS analysis of citrus orchards

In this study, the virome of citrus orchards in the main Spanish citrus growing area, the Valencian Community, was investigated by high-throughput sequencing (HTS). With this purpose 13 different orchards, representing the most common cultivars/rootstocks in the region, were selected in the three regional provinces: Valencia (8 orchards), Castellón (3 orchards) and Alicante (1 orchard). Total RNA isolated from leaf tissue of 20 citrus plants, both asymptomatic and showing viral-like symptoms of tree decline, leaf deformation and stunting was analyzed individually by HTS using Illumina technology. The bioinformatic analysis of the HTS data revealed the presence of several known viruses and viroids as well as unknown viruses (Table 2).

**Table 2.** Summary of the bioinformatic analysis results for each citrus sample analyzed by HTS in this study. Sample ID, the symptomatology observed in the plant, number of reads and contigs at different analysis steps and the viruses detected are shown.

| Sample ID | Symptomatology | Raw paired-end reads | Filtered reads <sup>a</sup> | De novo contigs | Viral-related contigs | Virus/viroid detected |
| --- | --- | --- | --- | --- | --- | --- |
| 211.1 | Asymptomatic | 28,332,696 | 6,598,226 | 23,444 | 104 | CTV genotype T30<br>CTV genotype RB |
| 211.2 | Asymptomatic | 28,802,804 | 6,309,699 | 18,826 | 28 | CTV genotype VT<br>*Botourmiaviridae* related |
| 216.2 | Asymptomatic | 23,897,672 | 2,517,443 | 16,405 | 15 | **CTV genotype T3**<br>CTV genotype T30<br>CVEV<br>*Botourmiaviridae* related<br>Mononegavirales related |
| 216.8 | Tree decline | 20,427,186 | 1,526,036 | 15,637 | 15 | CTV genotype T30<br>CVEV<br>*Solemoviridae* related<br>*Botourmiaviridae* related |
| 216.10 | Tree decline | 21,617,372 | 2,122,667 | 6,028 | 10 | **CTV genotype T3**<br>CTV genotype RB<br>*Botourmiaviridae* related |
| 249.1 | Tree decline | 25,874,721 | 1,584,934 | 4,086 | 22 | **CYVCV**<br>CTV genotype RB<br>CTV genotype VT<br>CLBV<br>CDVd<br>*Bromoviridae* related<br>Mononegavirales related |
| 256.5 | Stunting | 22,854,704 | 680,933 | 1,519 | 2 | CTV genotype VT<br>CTV genotype T30 |
| 260.5 | Tree decline | 20,246,082 | 142,692 | 2,032 | 12 | CTV genotype RB<br>CVEV<br>CDVd<br>CBLVd<br>HSVd<br>CEVd |
| 264.6 | Tree decline | 15,749,083 | 74,433 | 712 | 4 | CVEV<br>CTV genotype T30<br>*Tombusviridae* related |
| 267.6 | Tree decline | 24,997,805 | 620,234 | 941 | 18 | **CiVA**<br>CPsV<br>CTV genotype RB<br>CTV genotype VT<br>CTV genotype T30<br>CEVd<br>HSVd<br>CBLVd |
| 283.6 | Tree decline | 27,279,856 | 237,994 | 6,334 | 4 | **CTV genotype T3**<br>CTV genotype RB<br>CTV genotype VT<br>*Botourmiaviridae* related |
| 292 | Asymptomatic | 20,387,067 | 768,616 | 2,958 | 2 | CTV genotype RB |
| 298.1 | Leaf deformation | 27583083 | 1,079,915 | 3,368 | 10 | **CYVCV**<br>CTV genotype VT<br>CTV genotype T30<br>*Solemoviridae* related<br>Mononegavirales related |
| 314.1 | Asymptomatic | 24,551,571 | 246,559 | 5,572 | 19 | CTV genotype RB<br>CTV genotype VT<br>CTV genotype T30 |
| 314.2 | Asymptomatic | 22,831,078 | 175,283 | 4,076 | 14 | CTV genotype VT<br>CTV genotype RB |
| 314.3 | Asymptomatic | 26,247,367 | 418,402 | 2,350 | 17 | CTV genotype RB<br>CTV genotype VT |
| 314.4 | Asymptomatic | 24,683,230 | 60,566 | 6,737 | 4 | CTV genotype VT<br>CTV genotype T30<br>*Botourmiaviridae* related |
| 315.1 | Asymptomatic | 27,324,977 | 310,608 | 2,484 | 17 | CTV genotype T30<br>CTV genotype VT |
| 315.2 | Asymptomatic | 26,438,324 | 1,427,032 | 15,329 | 8 | CTV genotype RB<br>CTV genotype VT<br>*Botourmiaviridae* related |
| 315.3 | Asymptomatic | 26,891,097 | 158,103 | 408 | 6 | **CYVCV**<br>CTV unclassified genotype |

<sup>a</sup> Reads retained after adapter trimming, quality filtering, taxonomic filtering (viral and unclassified reads), and rRNA removal.

Among the known viruses detected in the HTS analysis, CYVCV, CiVA and CTV genotype T3 represent new relevant virome components related to important citrus diseases not previously reported in Spain. Other known viruses and viroids detected in the study had already been reported to infect citrus in the Spanish Mediterranean region, including T30, RB and VT genotypes of CTV, CLBV, CVEV, CPsV, CDVd, CBLVd, CEVd and HSVd. Moreover, several other viral-related sequences potentially representing the discovery of new unknown viruses were also found. Taken together, all these results point to the presence in citrus Spanish orchards of a much more complex virome than previously thought.

### Detection and molecular characterization of CYVCV in Spanish citrus orchards

HTS analysis of samples 249.1 and 298.1, corresponding to mandarin and sweet orange trees showing tree decline and leaf deformation symptoms, respectively, allowed the recovery of 2 large contigs of 7,530 and 7,462 nt with a sequence similarity of 97.59 % and 97.71 % respect to the CYVCV isolate KPMI (KT696513) from India. Further extension and curation steps resulted in the assembly of two almost full length CYVCV genomes of 7,590 nt, isolate IVIA 249.1 (PX970359) and isolate IVIA 298.1 (PX970360). BLAST analysis of these two genomes revealed a sequence similarity of 97.59 % and 97.60 %, as well as 97.44 % and 97.48 % with the isolates Y2 (MT951237) from Turkey and KPMI (KT696513) from India, respectively. A total of 4 small contigs ranging from 1,450 to 2,088 nt *de novo* assembled in sample 315.3, corresponding to an asymptomatic sweet orange tree, also showed sequence similarity by BLAST analysis with CYVCV. However, despite the detection of the virus, efforts to recover a full/almost full length CYVCV genomic sequence from this sample were unsuccessful.

Besides CYVCV, other known/unknown viruses/viroids were detected in these 3 samples: RB and VT genotypes of CTV, CLBV and CDVd in sample 249.1; T30 and VT genotypes of CTV in sample 298.1; and unclassified genotype of CTV in sample 315.3. In addition, sequences tentatively representing new members of the families *Bromoviridae* (sample 249.1), *Solemoviridae* (sample 298.1) as well as potential new member classified into Mononegavirales order (sample 298.1) were also detected.

The presence of CYVCV in samples 249.1 and 298.1 was confirmed by RT-PCR using primers previously reported in the literature for the amplification of two different genomic regions, a 487 nt fragment in the CP/NABP/3’ UTR region (Loconsole *et al*., 2012) and a 612 nt fragment in the CP region (Chen *et al*., 2014). Both regions were successfully amplified in these samples. Sanger sequencing of the amplicons confirmed 100 % the HTS determined sequence in all cases.

RT-PCR analysis using the primers described above was performed on 68 additional samples from the orchards prospected for HTS and other closely located orchards, including mandarin, sweet orange, lime and lemon trees. CYVCV was detected in 11 of the 68 samples analyzed. Among the positive samples, only 2 lime and 4 lemon trees showed leaf deformation and vein clearing symptoms compatible with CYVCV infection whereas 4 sweet orange CYVCV positive samples were symptomless, and 1 mandarin showed symptoms of stunting and tree decline. These results demonstrate the presence of CYVCV in Spanish citrus orchards and represent the first report of this pathogen in Spain.

A phylogenetic analysis conducted on 82 CYVCV complete genomes available in the databases, including the two full length Spanish genomes determined in this study revealed two main clusters separated with a high bootstrap support (72%) according to previous results (Sun and Yokomi, 2024). One cluster comprised the majority of Chinese CYVCV isolates also including South Korean isolates, whereas the other grouped the majority of isolates from the USA and isolates from India, Turkey and Italy. The Spanish isolates clustered within this latter group and formed a subcluster most closely related to one Turkish (MT951237) and one Indian isolate (KT696513) (Figure 1).

**Figure 1.**
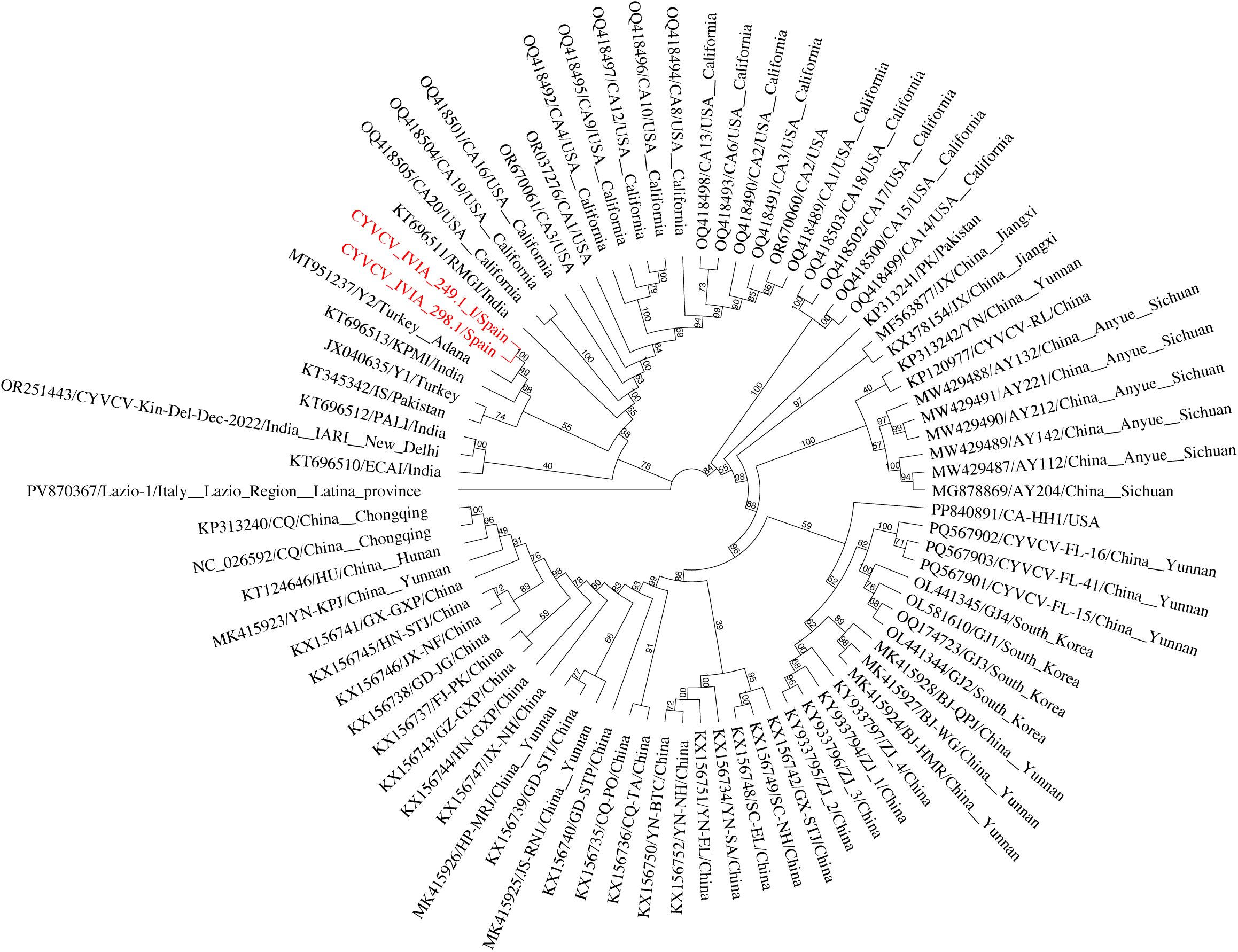
Full-length CYVCV genomic sequences maximum likelihood (ML) phylogenetic tree inferred using the General Time Reversible (GTR) +G +I model. The tree with the highest log likelihood is shown. Bootstrap support values obtained from 1000 replicates are indicated on the branches. A total of 82 complete CYVCV genome sequences were included in the analysis. Information on isolate identification, GenBank accession number and geographic origin is included. Spanish isolates are highlighted in red.

### Detection of CTV T3 genotype in Spanish citrus orchards

The bioinformatic analysis of HTS data from samples 216.2, an asymptomatic sweet orange tree, as well as 216.10 and 283.6, both sweet orange trees showing tree decline symptomatology, revealed the presence of 3 large contigs of 19,370, 13,675 and 19,161 nt, with a sequence similarity of 95.77%, 95.49 % and 95.86 with the CTV isolate T3-KB (MH051719.1) respectively, representing almost full length genomes of CTV T3 genotype. Extension and curation of these contigs allowed the recovery of 3 almost complete genomes of CTV genotype T3, isolate IVIA 216.2 (19,253 nt; PX970362), isolate IVIA 216.10 (19,253 nt; PX970361) and isolate IVIA 283.6 (19,185 nt). A phylogenetic analysis conducted on these 3 genomes confirmed their classification into CTV T3 genetic group. The neighbor-network reconstruction revealed that the Spanish CTV T3 isolates grouped clearly within the T3 genotype, clustering with isolates from Iran, South Africa, New Zealand and the USA, which represent the complete T3 genomes currently available in the database (Figure 2).

**Figure 2.**
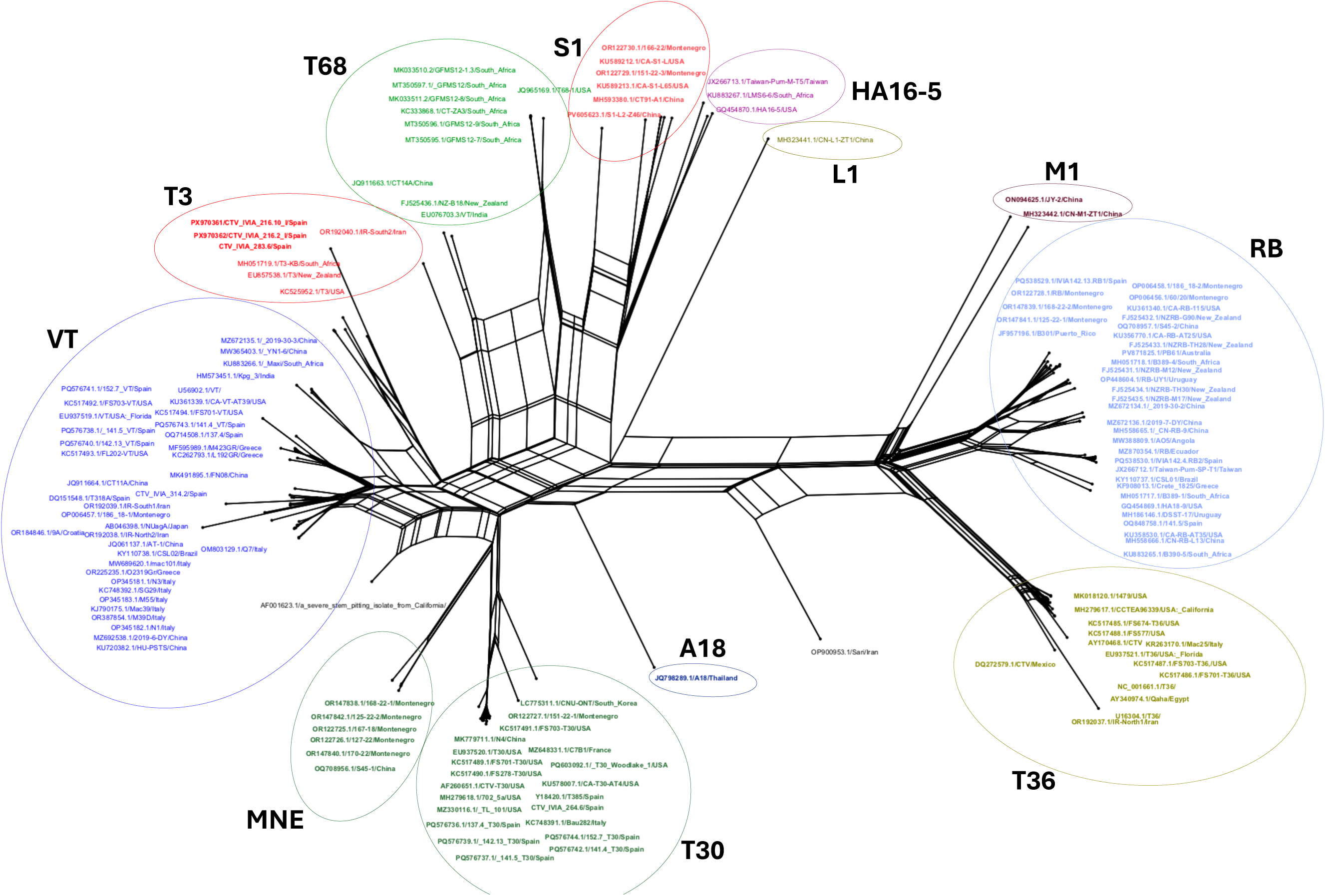
Neighbor network reconstruction of the complete genomes of citrus tristeza virus (CTV) based on 146 complete genome sequences. The analysis was performed using SplitsTree, applying the P-distance method to generate a pairwise distance matrix, followed by Neighbor Net inference. Genotype groups are indicated with colored circles. CTV isolates detected in this study are highlighted in bold within the T3 genotype.

**Figure 3.**
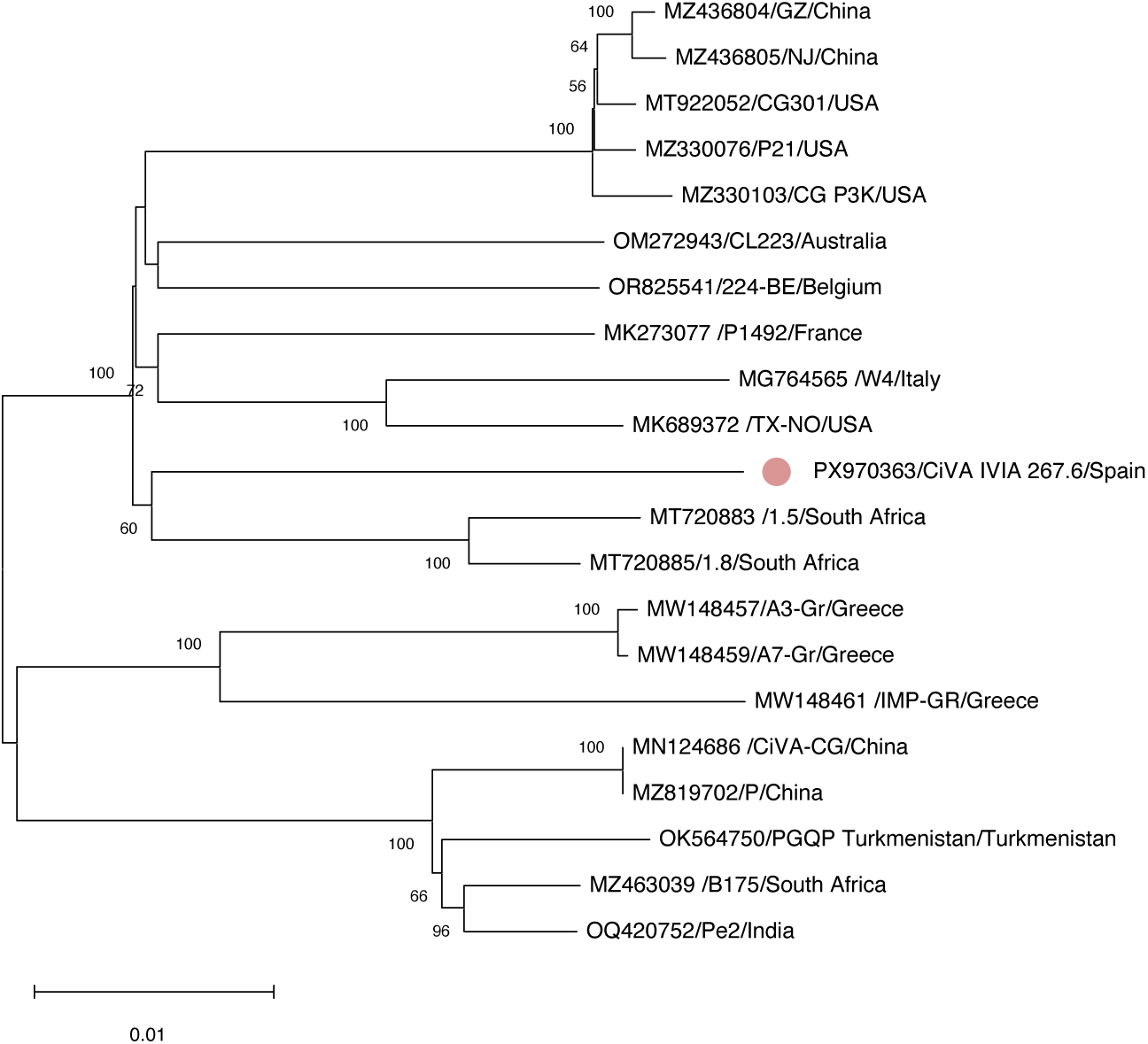
Full-length CiVA RNA 1 genomic sequences maximum likelihood (ML) phylogenetic tree inferred using the General Time Reversible (GTR) +G +I model. The tree with the highest log likelihood is shown. Bootstrap support values obtained from 1000 replicates are indicated on the branches. A total of 21 complete CiVA RNA 1 sequences were included in the analysis. Spanish isolates are highlighted with a colored dot.

Other viruses were detected in these CTV T3 infected trees: T30 genotype of CTV and CVEV in sample 216.2; RB genotype of CTV in sample 216.10; and RB and VT genotypes in sample 283.6. Moreover, viral-like sequences probably representing new uncharacterized viruses belonging to *Botourmiaviridae* (samples 216.2, 216.10 and 283.6) families, as well as Mononegavirales order (sample 216.2) were discovered.

### Characterization of the first Spanish CiVA genomic sequence

Among the viral-related contigs obtained in the bioinformatic analysis of the HTS data obtained from sample 267.6, a mandarin tree showing tree decline, 2 contigs of 4,931 and 1,592 nt showing 95.81 % and 95.91 % sequence similarity with RNA 1 of the CiVA isolates 224-BE (OR825541, Belgium) and 1.8 (MT720885, South Africa) and 1 contig of 2,729 nt showed sequence similarity of 96.12 % with RNA 2 of the CiVa isolate CL223 (OM272944, Australia). Extension and curation of the contigs resulted in the recovery of almost complete genomic regions of RNA 1 (6,646 nt) and RNA 2 (2,729 nt) of the first identified Spanish genome of this viral species. RNA 1 sequence (IVIA 267.6 RNA 1, PX970363) showed 95.73 % nucleotide identity with the pear isolate 224-BE (OR825541) from Belgium. RNA 2 (IVIA 267.6 RNA2, PX970364) shared 96.12 % sequence similarity with the isolate CL223 (OM272944) from Australia.

CPsV, T30, RB and VT genotypes of CTV, and the viroids CEVd, HSVd and CBLVd were also detected in sample 267.6 by HTS.

In order to confirm the presence of CiVA in this sample two different genomic regions were tested by RT-PCR using primers previously described amplifying 703 nt in the RdRp region of RNA 1 (Fontdevila *et al*., 2025) and 285 nt located in the 5’UTR/MP region of RNA 2 (Bester *et al*., 2021). RT-PCR amplification was successfully achieved for RNA 2 detection, and Sanger sequencing of the obtained amplicon confirmed 100 % the HTS recovered sequence of CiVA RNA 2. Concerning RNA1, unsuccessful efforts were carried out to amplify the RdRp region mentioned above. Consequently, a new set of primers targeted to a different RdRp region of 390 nt in RNA 1 were designed (listed on Table 2). Using this primer set the RNA 1 CiVA HTS recovered sequence could be confirmed 100% by RT-PCR amplification and Sanger sequencing.

RT-PCR analysis using primers designed by Bester *et al*., 2021 on CiVA RNA 2 was performed on the 68 additional citrus samples described above. This analysis revealed the presence of this virus in 8 of the 68 samples, corresponding to 2 mandarin trees showing tree decline, 4 asymptomatic orange trees and 2 asymptomatic mandarin trees. It is important to note that fruits were not evaluated during the survey. Therefore, the fruit symptomatology potentially associated to the presence of CiVA in Spanish orchards remains to be studied.

Despite the already stablished occurrence of imprietatura disease in Spain, our finding represents the first characterization of CiVA genomic sequence in Spanish citrus orchards.

A phylogenetic study on 20 CiVA RNA1 complete sequences available in the databases and the Spanish sequence here obtained identified three main clusters (C1, C2 and C3) with strong bootstrap support (100%). The Spanish CiVA isolate grouped within cluster C1, which comprises isolates from multiple countries, showing the closest phylogenetic relationship with two South African isolates (MT720883 and MT720885).

## Discussion

This study provides an updated and more comprehensive view of the citrus virome in Spain revealing the complexity of the Spanish citrus virome that includes in addition to well-known regulated pathogens other new and recently described viruses. These findings highlight the value of systematic virome-based surveillance strategies in citrus-growing areas to update and reinforce routinary targeted diagnostic approaches.

The detection of CYVCV in the surveyed orchards adds a further element to the changing citrus virome in the EPPO region. CYVCV is a potexvirus infecting a broad range of citrus species and is associated with vein clearing and chlorosis, leaf deformation and, under severe conditions, reductions in tree vigour and fruit quality (Liu *et al*., 2020; Abrahamian *et al*., 2024). Although the virus has been known for decades in parts of Asia and the Middle East, recent detections indicate an ongoing geographical expansion and an increasing likelihood of emergence into new citrus-producing areas (Jin *et al*., 2024; Cinque *et al*., 2024).

CYVCV can be efficiently transmitted through vegetative propagation, including grafting and the movement of infected budwood, which represents the main pathway for long-distance spread. Local dissemination is mainly driven by insect-mediated transmission and, to a lesser extent, by mechanical transmission (Zhang *et al*., 2018a). Experimental evidence has shown that several hemipteran species are competent vectors of CYVCV, including aphids such as *Aphis aurantii*, *Aphis gossypii* and *Aphis spiraecola*, the whitefly *Dialeurodes citri* (Aleyrodidae), and the psyllid *Diaphorina citri* (Psyllidae) (Önelge et al., 2011; Zhang *et al*., 2018b; Zhang *et al*., 2019; Liu *et al*., 2020; Maghsoudi *et al*., 2023).

The host range of CYVCV include most commercially important citrus species, and infections can be latent in several hosts, with symptom expression depending on cultivar, physiological status of the plant and environmental conditions. Symptom severity is often higher in lemon and sour orange, whereas infections in other citrus species may remain asymptomatic or show transient symptoms that are masked under high temperatures or in mature leaves (Iftikhar *et al*., 2010; Zhou *et al*., 2017.). Recently, CYVCV has also been described as the causal agent of spring shoot leaf curl disease in mandarin (Liu *et al*., 2025). Its variability in symptom expression complicates field diagnosis and favours unnoticed circulation of the virus in orchards and nurseries.

From a risk perspective, the recent detection of CYVCV within the EPPO region, specifically in Italy (Cinque *et al*., 2024) together with its presence in major citrus-producing countries outside Europe, indicates that the virus is no longer geographically confined and may continue to expand its range (Jin *et al*., 2024; Cinque *et al*., 2024). The potential impact in Mediterranean citrus systems is expected to depend on interactions among virus isolates, host species and local agronomic and climatic conditions. Experimental and field data indicate that CYVCV infection can affect physiological processes related to photosynthesis and nutrient transport, providing a mechanistic basis for yield and quality losses in susceptible hosts. In this context, the identification of CYVCV by HTS in our study is epidemiologically relevant, as it reveals the presence of an emerging virus that may have remained undetected by symptom-based surveys, particularly in asymptomatic hosts or outside optimal phenological stage. The ongoing pest risk analysis within the EPPO framework reflects the increasing recognition of CYVCV as a phytosanitary concern. Our results provide field-based evidence supporting the inclusion of CYVCV in structured surveillance programs and highlight the value of HTS to detect early introductions before widespread establishment occurs.

The detection of a new CTV genotype, corresponding to the T3 genotype, represents another relevant epidemiological finding. This genotype had not been previously reported in Spain. Although CTV is a long-established pathogen in Spanish citrus orchards, the presence of additional genotypes expands the known genetic diversity of the virus in the country and raises questions about its origin, pathways of introduction and potential biological behaviour under local and Mediterranean conditions. Given the well-documented differences in pathogenicity, transmission efficiency and interactions among CTV genotypes, continuous molecular surveillance is required to monitor potential changes in population structure and associated risks.

The current classification of CTV relies on viral genotype rather than on the type or severity of symptoms observed in infected trees (Dawson *et al*., 2015). At present, twelve genotypes or phylogenetic groups are recognized: T30, T36, T3, T68, VT, RB, HA16-5, S1, M1, L1, A18 and MNE (Harper, 2013; Yokomi *et al*., 2018; Bester *et al*., 2021; Zindović *et al*., 2024). The traditional distinction between “mild” and “severe” CTV isolates is therefore of limited value, as isolates within the same genotype may be associated with very different symptomatology. For example, isolates belonging to T36 and VT have been linked to both mild infections and severe syndromes such as tristeza disease, stem pitting and seedling yellows (Varveri *et al*., 2015; EPPO, 2023). In contrast, T30, S1 and RB isolates are more often associated with latent infections or mild symptoms, whereas T68 and T3 are more frequently linked to severe disease expression (Albiach-Marti, 2013; Yokomi *et al*., 2018; PM 7/32 (2) EPPO, 2023).

Under field conditions, mixed infections involving several CTV genotypes within the same tree are common and have been associated with a wide range of symptom expression depending on host species and cultivar. Interestingly, among all the CTV genetic diversity observed worldwide, only a subset of this global diversity has been historically detected in the EPPO region, where some phylogenetic groups remain absent and others have a limited geographical distribution. As a result, CTV populations circulating in Europe represent only a fraction of the worldwide diversity reported for this virus (EFSA PLH Panel, 2017). Within this framework, the recent description of RB genotype CTV in Spain (Ruiz-García and Olmos, 2025) and the identification of the T3 genotype in Spain reported in this study are epidemiologically relevant, as they reflect a further increase in the genetic heterogeneity of CTV in the region. The introduction and establishment of genotypes that have been uncommon or absent in Europe may alter the current population structure of CTV and potentially modify disease expression and epidemiological behaviour. Thus, the epidemiology of CTV further supports the need for continuous surveillance, reinforcing the need to move beyond virus species-level detection and to incorporate phylogenetic characterization into surveillance programs. HTS-based approaches are particularly well-suited for this purpose, as they allow the simultaneous identification of known and emerging CTV genotypes and provide a robust framework for monitoring changes in virus populations over time. This level of resolution is essential to anticipate potential shifts in disease impact and to support evidence-based updates of phytosanitary strategies.

The detection of CiVA in Spanish orchards is relevant in the context of citrus diseases with previously unclear etiology. Since its initial description in citrus (Navarro *et al*., 2018), CiVA has been reported in different citrus-growing regions (Bester *et al*., 2021) and associated with citrus impietratura disease, supported by several complementary observations (Beris *et al*., 2021). Although impietratura disease has been reported in Spain (EPPO Global Database), CiVA had not been reported to date in our territory nor Spanish CiVA genomic sequences had been described. In the present study, CiVA was detected in citrus trees sampled at a time when no fruits were available. For this reason, the presence of impietratura symptoms could not be evaluated in the analyzed material. Further surveys during the fruiting period will be required to determine whether CiVA-infected trees in the surveyed orchards develop typical impietratura symptoms under local conditions. Nevertheless, beyond its association with impietratura, the detection of CiVA in commercial orchards has phytosanitary implications, as this virus is currently listed as a regulated pest that should be absent from certified citrus propagation material. The availability of HTS-based molecular genomic information supports the development of practical detection tools to support certification schemes and to improve surveillance of citrus-growing areas. Overall, the detection of CiVA in the analyzed orchards also extends the geographical and epidemiological context of this virus.

Taken together, the detection in Spanish orchards of CYVCV, alongside the identification of a new CTV genotypes as well as CiVA, illustrates the dynamic nature of the citrus virome in the EPPO region. These findings support a shift towards risk-based surveillance strategies that should integrate HTS with targeted field surveys and reinforce the need to periodically re-evaluate the list of regulated citrus viruses in light of emerging epidemiological evidence and coordinated international risk assessments.

Moreover, beyond well-characterised citrus viruses and viroids, the virome analysis revealed the presence of sequences related to several viral families, including *Botourmiaviridae*, *Solemoviridae*, *Bromoviridae* and members of the order *Mononegavirales*, that might account for citrus unknown viruses. The biological nature, host range and epidemiological relevance of these virus-like sequences in citrus should be investigated. The identification of these putative novel or poorly characterized viral components further illustrates the power of HTS-based approaches to uncover hidden viral diversity in agroecosystems.

In conclusion, this study provides the first evidence of the presence of CYVCV, CTV T3 genotype and CiVA in Spain, placing these findings within a broader context of an increasingly complex and dynamic citrus virome in Spain. The detection of emerging and previously unreported viral agents in commercial orchards support the inclusion of virome-level analyses in phytosanitary surveillance and certification schemes, particularly for the assessment of propagation material.

---

**Supplementary Table S1.**
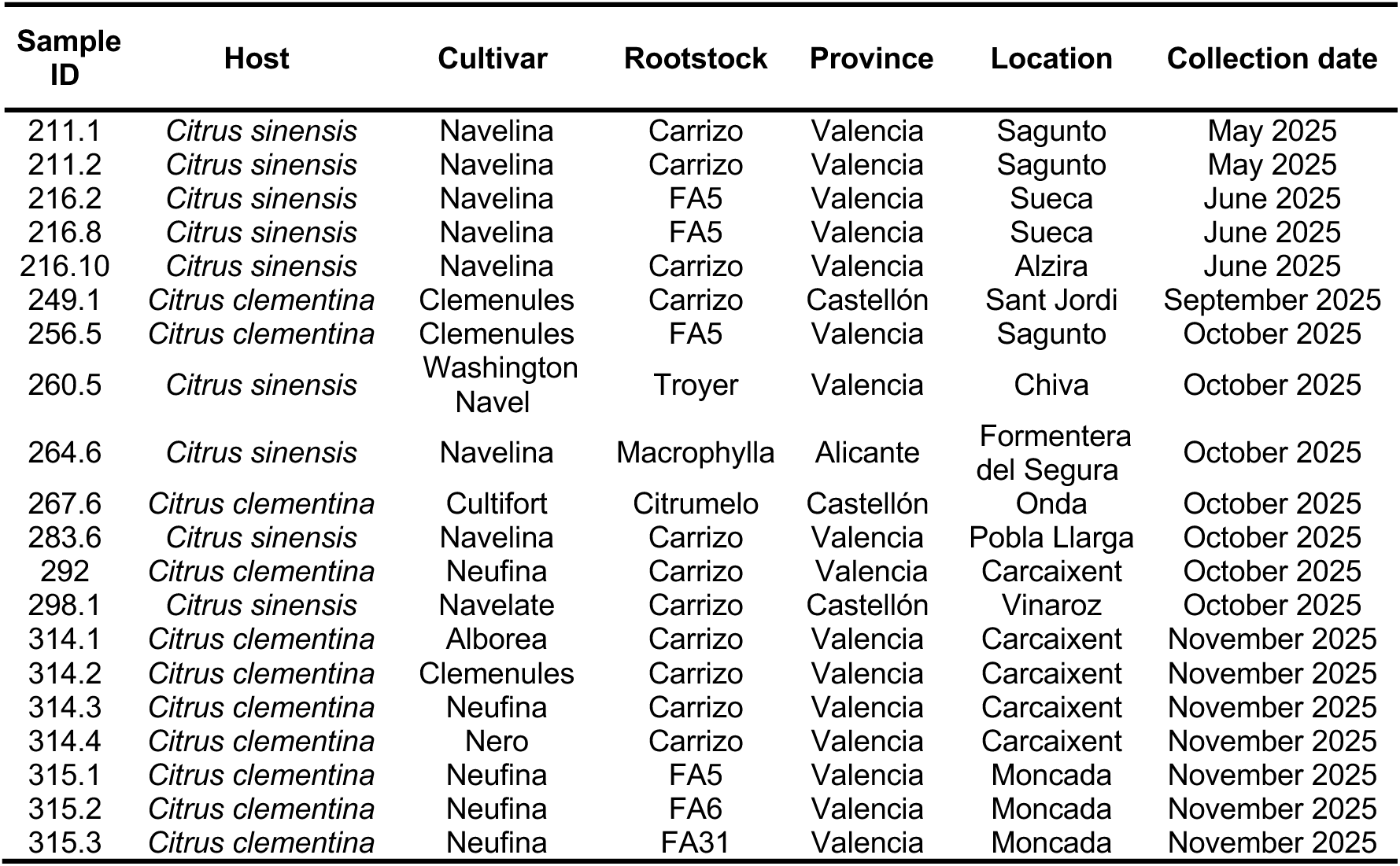
Description of citrus samples analyzed in this study by HTS, including sample identification code, host species, cultivar, rootstock, province and locality of origin and collection date.

| Sample ID | Host | Cultivar | Rootstock | Province | Location | Collection date |
| --- | --- | --- | --- | --- | --- | --- |
| 211.1 | *Citrus sinensis* | Navelina | Carrizo | Valencia | Sagunto | May 2025 |
| 211.2 | *Citrus sinensis* | Navelina | Carrizo | Valencia | Sagunto | May 2025 |
| 216.2 | *Citrus sinensis* | Navelina | FA5 | Valencia | Sueca | June 2025 |
| 216.8 | *Citrus sinensis* | Navelina | FA5 | Valencia | Sueca | June 2025 |
| 216.10 | *Citrus sinensis* | Navelina | Carrizo | Valencia | Alzira | June 2025 |
| 249.1 | *Citrus clementina* | Clemenules | Carrizo | Castellón | Sant Jordi | September 2025 |
| 256.5 | *Citrus clementina* | Clemenules | FA5 | Valencia | Sagunto | October 2025 |
| 260.5 | *Citrus sinensis* | Washington Navel | Troyer | Valencia | Chiva | October 2025 |
| 264.6 | *Citrus sinensis* | Navelina | Macrophylla | Alicante | Formentera del Segura | October 2025 |
| 267.6 | *Citrus clementina* | Cultifort | Citrumelo | Castellón | Onda | October 2025 |
| 283.6 | *Citrus sinensis* | Navelina | Carrizo | Valencia | Pobla Llarga | October 2025 |
| 292 | *Citrus clementina* | Neufina | Carrizo | Valencia | Carcaixent | October 2025 |
| 298.1 | *Citrus sinensis* | Navelate | Carrizo | Castellón | Vinaroz | October 2025 |
| 314.1 | *Citrus clementina* | Alborea | Carrizo | Valencia | Carcaixent | November 2025 |
| 314.2 | *Citrus clementina* | Clemenules | Carrizo | Valencia | Carcaixent | November 2025 |
| 314.3 | *Citrus clementina* | Neufina | Carrizo | Valencia | Carcaixent | November 2025 |
| 314.4 | *Citrus clementina* | Nero | Carrizo | Valencia | Carcaixent | November 2025 |
| 315.1 | *Citrus clementina* | Neufina | FA5 | Valencia | Moncada | November 2025 |
| 315.2 | *Citrus clementina* | Neufina | FA6 | Valencia | Moncada | November 2025 |
| 315.3 | *Citrus clementina* | Neufina | FA31 | Valencia | Moncada | November 2025 |

**Supplementary Table S2.** Description of citrus samples analyzed by RT-PCR in this study for the detection of CYVCV and CiVA, including sample identification code, host species, cultivar, rootstock, province and locality of origin and virus/es detected.

| Sample ID | Host | Cultivar | Rootstock | Province | Location | Virus/es Detected |
| --- | --- | --- | --- | --- | --- | --- |
| 216.1 | *Citrus sinensis* | Navelina | FA5 | Valencia | Sueca | - |
| 216.3 | *Citrus sinensis* | Navelina | FA5 | Valencia | Sueca | - |
| 216.4 | *Citrus sinensis* | Navelina | FA5 | Valencia | Sueca | - |
| 249.2 | *Citrus clementina* | Clemenules | Carrizo | Castellón | Sant Jordi | - |
| 249.3 | *Citrus clementina* | Clemenules | Carrizo | Castellón | Sant Jordi | - |
| 249.4 | *Citrus clementina* | Clemenules | Carrizo | Castellón | Sant Jordi | - |
| 249.5 | *Citrus clementina* | Clemenules | Carrizo | Castellón | Sant Jordi | - |
| 249.6 | *Citrus clementina* | Clemenules | Carrizo | Castellón | Sant Jordi | - |
| 256.1 | *Citrus clementina* | Clemenules | FA5 | Valencia | Sagunto | - |
| 256.2 | *Citrus clementina* | Clemenules | FA5 | Valencia | Sagunto | - |
| 256.3 | *Citrus clementina* | Clemenules | FA5 | Valencia | Sagunto | - |
| 256.4 | *Citrus clementina* | Clemenules | FA5 | Valencia | Sagunto | - |
| 256.6 | *Citrus clementina* | Clemenules | FA5 | Valencia | Sagunto | - |
| 260.1 | *Citrus sinensis* | Whashington Navel | Troyer | Valencia | Chiva | - |
| 260.2 | *Citrus sinensis* | Whashington Navel | Troyer | Valencia | Chiva | - |
| 260.3 | *Citrus sinensis* | Whashington Navel | Troyer | Valencia | Chiva | - |
| 260.4 | *Citrus sinensis* | Whashington Navel | Troyer | Valencia | Chiva | - |
| 260.6 | *Citrus sinensis* | Whashington Navel | Troyer | Valencia | Chiva | - |
| 264.1 | *Citrus sinensis* | Navelina | Macrophylla | Alicante | Formentera del Segura | - |
| 264.2 | *Citrus sinensis* | Navelina | Macrophylla | Alicante | Formentera del Segura | CiVA |
| 264.3 | *Citrus sinensis* | Navelina | Macrophylla | Alicante | Formentera del Segura | CiVA |
| 264.4 | *Citrus sinensis* | Navelina | Macrophylla | Alicante | Formentera del Segura | - |
| 264.5 | *Citrus sinensis* | Navelina | Macrophylla | Alicante | Formentera del Segura | - |
| 267.1 | *Citrus clementina* | Cultifort | Citrumelo | Castellón | Onda | - |
| 267.2 | *Citrus clementina* | Cultifort | Citrumelo | Castellón | Onda | - |
| 267.3 | *Citrus clementina* | Cultifort | Citrumelo | Castellón | Onda | - |
| 267.4 | *Citrus clementina* | Cultifort | Citrumelo | Castellón | Onda | CiVA |
| 267.5 | *Citrus clementina* | Cultifort | Citrumelo | Castellón | Onda | - |
| 274.1 | *Citrus clementina* | Clemenules | Carrizo | Castellón | Villareal | - |
| 274.2 | *Citrus clementina* | Clemenules | Carrizo | Castellón | Villareal | CiVA |
| 274.3 | *Citrus clementina* | Clemenules | Carrizo | Castellón | Villareal | - |
| 274.4 | *Citrus clementina* | Clemenules | Carrizo | Castellón | Villareal | - |
| 274.5 | *Citrus clementina* | Clemenules | Carrizo | Castellón | Villareal | - |
| 274.6 | *Citrus clementina* | Clemenules | Carrizo | Castellón | Villareal | - |
| 278.1 | *Citrus sinensis* | Navelina | Carrizo | Valencia | Carcaixent | CYVCV |
| 278.2 | *Citrus sinensis* | Navelina | Carrizo | Valencia | Carcaixent | - |
| 278.3 | *Citrus sinensis* | Navelina | Carrizo | Valencia | Carcaixent | CYVCV/CiVA |
| 278.4 | *Citrus sinensis* | Navelina | Carrizo | Valencia | Carcaixent | CYVCV |
| 278.5 | *Citrus sinensis* | Navelina | Carrizo | Valencia | Carcaixent | CYVCV |
| 278.6 | *Citrus sinensis* | Navelina | Carrizo | Valencia | Carcaixent | CiVA |
| 283.1 | *Citrus sinensis* | Navelina | Carrizo | Valencia | La Pobla Llarga | - |
| 283.2 | *Citrus sinensis* | Navelina | Carrizo | Valencia | La Pobla Llarga | - |
| 283.3 | *Citrus sinensis* | Navelina | Carrizo | Valencia | La Pobla Llarga | - |
| 283.4 | *Citrus sinensis* | Navelina | Carrizo | Valencia | La Pobla Llarga | - |
| 283.5 | *Citrus sinensis* | Navelina | Carrizo | Valencia | La Pobla Llarga | - |
| 287.1 | *Citrus clementina* | Clemenules | Citrumelo | Castellón | Castellón de la Plana | - |
| 287.2 | *Citrus clementina* | Clemenules | Citrumelo | Castellón | Castellón de la Plana | - |
| 287.3 | *Citrus clementina* | Clemenules | Citrumelo | Castellón | Castellón de la Plana | - |
| 287.4 | *Citrus clementina* | Clemenules | Citrumelo | Castellón | Castellón de la Plana | CiVA |
| 287.5 | *Citrus clementina* | Clemenules | Citrumelo | Castellón | Castellón de la Plana | - |
| 287.6 | *Citrus clementina* | Clemenules | Citrumelo | Castellón | Castellón de la Plana | - |
| 291.1 | *Citrus clementina* | Clemenules | Carrizo | Castellón | Betxí | CiVA |
| 291.2 | *Citrus clementina* | Clemenules | Carrizo | Castellón | Betxí | - |
| 291.3 | *Citrus clementina* | Clemenules | Carrizo | Castellón | Betxí | - |
| 291.4 | *Citrus clementina* | Clemenules | Carrizo | Castellón | Betxí | - |

| **Sample ID** | **Host** | **Cultivar** | **Rootstock** | **Province** | **Location** | **Virus/es Detected** |
| --- | --- | --- | --- | --- | --- | --- |
| **291.5** | *Citrus clementina* | Clemenules | Carrizo | Castellón | Betxí | CYVCV |
| **291.6** | *Citrus clementina* | Clemenules | Carrizo | Castellón | Betxí | - |
| **298.2** | *Citrus sinensis* | Navelate | Carrizo | Castellón | Vinaroz | - |
| **298.3** | *Citrus sinensis* | Navelate | Carrizo | Castellón | Vinaroz | - |
| **298.4** | *Citrus sinensis* | Navelate | Carrizo | Castellón | Vinaroz | - |
| **298.5** | *Citrus sinensis* | Navelate | Carrizo | Castellón | Vinaroz | - |
| **298.6** | *Citrus sinensis* | Navelate | Carrizo | Castellón | Vinaroz | - |
| **344.1** | *Citrus limon* | Unknown | Unknown | Castellón | Benicarló | CYVCV |
| **344.3** | *Citrus limon* | Unknown | Unknown | Castellón | Benicarló | CYVCV |
| **344.4** | *Citrus latifolia* | Unknown | Unknown | Castellón | Vinaroz | CYVCV |
| **344.5** | *Citrus limon* | Unknown | Unknown | Castellón | Vinaroz | CYVCV |
| **344.7** | *Citrus limon* | Unknown | Unknown | Castellón | Vinaroz | CYVCV |
| **344.8** | *Citrus latifolia* | Unknown | Unknown | Castellón | Vinaroz | CYVCV |

